# Impaired refinement of kinematic variability in Huntington disease mice on an automated home-cage forelimb motor task

**DOI:** 10.1101/2021.01.28.428530

**Authors:** Cameron L. Woodard, Marja D. Sepers, Lynn A. Raymond

## Abstract

The effective development of novel therapies in mouse models of neurological disorders relies on behavioural assessments that provide accurate read-outs of neuronal dysfunction and/or degeneration. We designed an automated behavioural testing system (‘PiPaw’) which integrates an operant lever-pulling task directly into the mouse home-cage. This task is accessible to group-housed mice 24-hours per day, enabling high-throughput longitudinal analysis of forelimb motor learning. Moreover, this design eliminates the need for exposure to novel environments and minimizes experimenter interaction, significantly reducing two of the largest stressors associated with animal behaviour. Mice improved their performance of this task over one week of testing by reducing inter-trial variability of reward-related kinematic parameters (pull amplitude or peak velocity). In addition, mice displayed short-term improvements in reward rate, and a concomitant decrease in movement variability, over the course of brief (<10 minutes) bouts of task engagement. We used this system to assess motor learning in mouse models of the inherited neurodegenerative disorder, Huntington disease (HD). Despite having no baseline differences in task performance, Q175-FDN HD mice were unable to modulate the variability of their movements in order to increase reward on either short or long timescales. Task training was associated with a decrease in the amplitude of spontaneous excitatory activity recorded from striatal medium spiny neurons in the hemisphere contralateral to the trained forelimb in wildtype mice; however, no such changes were observed in Q175-FDN mice. This behavioural screening platform should prove useful for preclinical drug trials towards improved treatments in HD and other neurological disorders.

**Significance Statement:** In order to develop effective therapies for neurological disorders such as Huntington disease (HD), it’s important to be able to accurately and reliably assess the behaviour of mouse models of these conditions. Moreover, these behavioural assessments should provide an accurate readout of underlying neuronal dysfunction and/or degeneration. In this paper, we employed an automated behavioural testing system to assess motor learning in mice within their home-cage. Using this system, we were able to study motor abnormalities in HD mice with an unprecedented level of detail, and identified a specific behavioural deficit associated with an underlying impairment in striatal neuronal plasticity. These results validate the usefulness of this system for assessing behaviour in mouse models of HD and other neurological disorders.

## Introduction

Huntington disease (HD) is a dominantly inherited neurodegenerative disorder caused by a CAG-repeat expansion in the gene encoding the protein huntingtin (HTT). HD is characterized by selective neurodegeneration which most prominently affects the striatum, a key structure of the basal ganglia involved in motor function and action selection (Graybiel and Grafton, 2015; Klaus et al., 2019). In those affected by HD, degeneration of striatal medium spiny neurons (MSNs) results in progressive motor dysfunction, and together with cortical degeneration, also cognitive impairment and neuropsychiatric symptoms (Bates et al., 2015). Characteristic motor symptoms of HD include chorea, bradykinesia, rigidity and difficulties with balance and gait (McColgan and Tabrizi, 2018). However, individuals affected by HD also have deficits in their ability to learn and control voluntary movements, some of which appear years before the onset of overt motor dysfunction (Bonfiglioli et al., 1998; Smith et al., 2000; Klein et al., 2011; Shabbott et al., 2013).

Several transgenic and knock-in mouse models of HD have been generated which display a similar pattern of neurodegeneration as that seen in HD patients (Pouladi et al., 2013). These mice also have diverse behavioural alterations, although the specifics of their phenotype and the timing of motor deficits vary widely between models. Unfortunately, due to the complex phenotype displayed by HD mice, the read-outs given by some commonly used behavioural paradigms are difficult to interpret. For example, some transgenic models of HD have increased bodyweight (Pouladi et al., 2010), a known confound for assessments of full-body motor coordination such as the rotarod test (McFadyen et al., 2003). Some HD models also display anxiety-like behaviour on assessments of approach-avoidance conflict (e.g. the elevated plus maze) (Abada et al., 2013; Glangetas et al., 2020). As many behavioural assessments involve exposure to novel and brightly lit areas, this phenotype could systematically bias performance on these tests. Furthermore, mouse handling and olfactory exposure to experimenters have been reported to cause stress responses in mice (Balcombe et al., 2004; Sorge et al., 2014). These physiological responses may influence baseline behavioural abnormalities in HD mice, masking or enhancing genotype differences and confounding pre-clinical research.

An alternative to traditional behavioural testing paradigms is to integrate an operant task into the mouse home-cage. By assessing mice on a self-paced task within their home environment, exposure of mice to handling and other stressors is significantly reduced, and animals are given greater control over their interactions with the task. This allows learning to progress in a more naturalistic and self-directed fashion and facilitates collection of longitudinal behavioural datasets. Notably, several studies have integrated forelimb motor tasks into the rodent home-cage (Poddar et al., 2013; Woodard et al., 2017; Silasi et al., 2018; Bollu et al., 2019; Salameh et al., 2020). Moreover, given reported deficits in voluntary arm and hand movements in HD patients, tests of forelimb motor learning may provide a measure of motor function in HD mice less biased by confounds such as body weight. Indeed, two previous studies have found deficits on reaching and lever manipulation tasks in HD model mice (Woodard et al., 2017; Glangetas et al., 2020), encouraging further investigation.

Here, we employed an automated home-cage system (‘PiPaw’) to assess forelimb motor learning in group-housed mice. Mice tested in this system learned to grasp a one-axis lever with their forelimb and perform pulls of a specific amplitude in order to receive water drops. We found that wildtype mice rapidly acquired this task and improved their performance over time by reducing the variability of reward-related kinematic parameters. In contrast, Q175-FDN HD mice were unable to modulate the variability of their movements in order to increase reward on either short or long timescales. Furthermore, we found that reduced variability was associated with a decrease in the average amplitude of spontaneous activity of contralateral striatal MSNs in wildtype, but not Q175-FDN mice, suggesting that the observed motor learning deficits are related to impaired striatal neuronal plasticity.

## Materials and Methods

### Animals

All procedures were carried out in accordance with the Canadian Council on Animal Care and approved by the University of British Columbia Committee on Animal Care (protocol A19-0076). Experiments were conducted using 2- to 3-month-old FVB/N mice, as well as 10- to 11-month-old heterozygous YAC128 transgenic mice (line 53; Slow et al., 2003) and heterozygous Q175-FDN knock-in mice (Southwell et al., 2016), both on the FVB/N genetic background. All mice were male, and wildtype littermates of transgenic and knock-in mice were used as controls. Animals were housed on a 12/12 h light/dark cycle in a temperature and humidity-controlled room and were provided with standard environmental enrichment within the cage (bedding, hut, PVC tube) throughout testing. Animal tissue was collected via ear clipping at weaning, and DNA extraction and PCR analysis was subsequently used to determine genotype. In order to enable automated identification of group-housed mice during behavioural testing, glass RFID capsules (Sparkfun SEN-09416) were subcutaneously implanted in the upper thoracic torso as previously described (Woodard et al., 2020). Mice were handled by the experimenter on at least two occasions for 2-3 minutes per animal prior to any surgery or behavioural testing.

### PiPaw task

#### Hardware and software

PiPaw was developed based on the design of an earlier home-cage behavioural testing system (Woodard et al., 2017; Silasi et al., 2018). An opening was created on one side of a regular mouse home-cage to allow mice to access an attached 3D-printed chamber (the ‘testing module’) (Fig. 1a). At the opposite end of the module from the entrance, a nose-poke port accessed a water spout, which delivered drops using a gravity-fed valve-based system (Murphy et al., 2016). An RFID antenna and reader (Sparkfun SEN-11828) were inset into the ceiling in order to detect and identify animals. On the right wall of the chamber, adjacent to and slightly below the nose poke port, a lever extended 1.5 cm into the chamber. This lever was moveable on a horizontal axis with a range of 30° (~1 cm at the end of the lever) and was positioned such that the mouse’s right forelimb would naturally rest on it when accessing the nose-poke port. Across from the lever, a small ledge allowed the mouse to support themselves on their left forelimb while nose-poking and grasping the lever.

**Figure 1:**
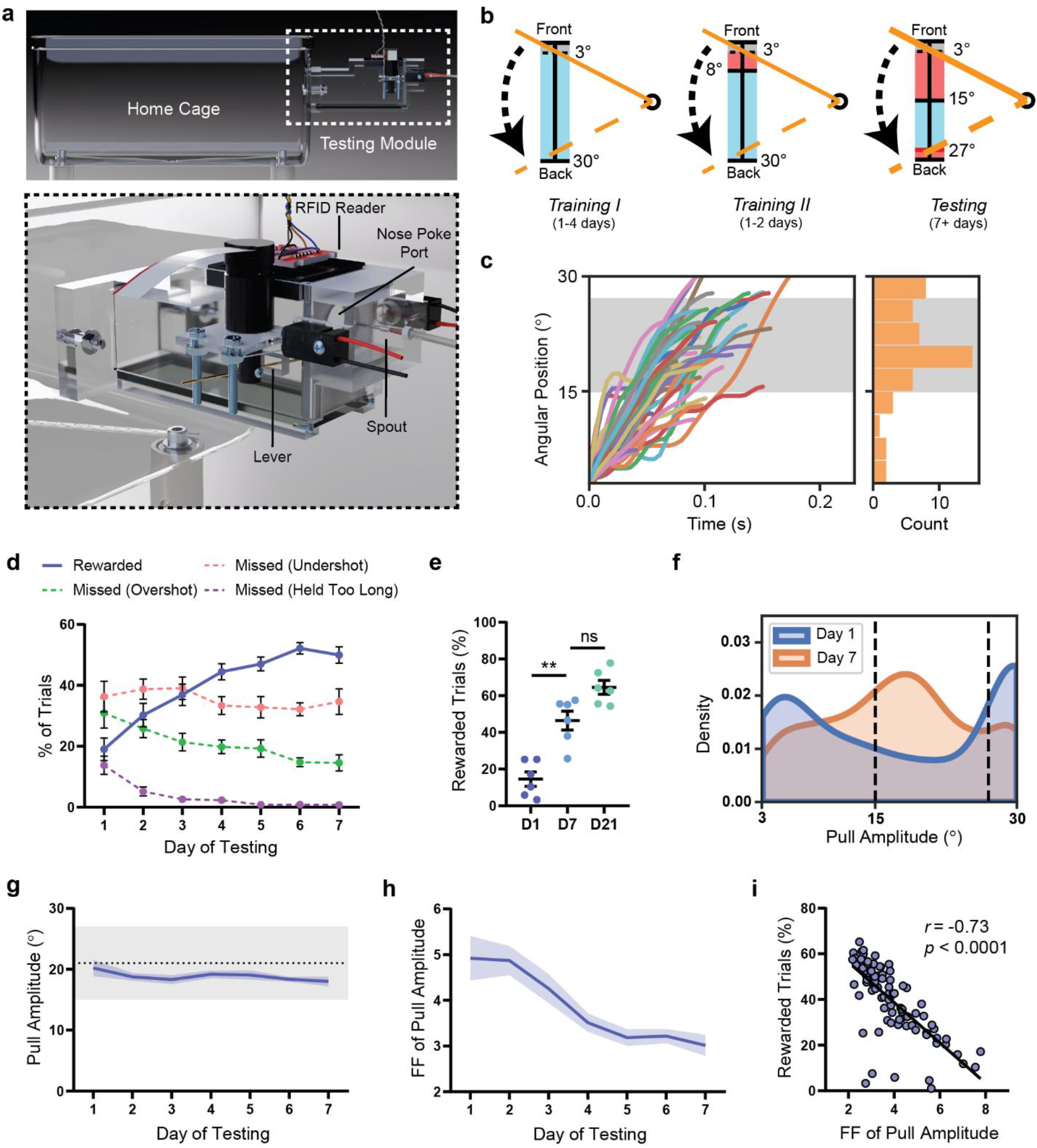
WT mice improve their performance of the PiPaw task by decreasing the inter-trial variability of pull amplitude. **(a)** A small testing module was attached to the mouse home-cage containing a lever, a nose-poke port and a spout which delivered water drop rewards. An RFID reader would detect and identify animals upon entrance into the chamber and load the appropriate testing parameters. In order to perform a trial, the mouse was required to nose-poke and simultaneously pull the lever using their right forelimb. **(b)** The lever was held in its starting position at the front of the lever position range by the motor until a trial was initiated. In all testing phases, the mouse initiated a trial by pulling the lever backwards out of the threshold range (0-3°, shown in grey). In Training I, a water drop reward was given for all trials, regardless of pull amplitude, as long as the lever was returned back to the start position before the trial time limit (2 s). In Training II, the trial was rewarded only if the amplitude of the pull (i.e. maximum position of the lever during the trial) was greater than 8°. In the Testing phase, the rewarded pull amplitude range narrowed to between 15° and 27°. If the amplitude of the pull was less than 15° (undershot) or greater than 27° (overshot), the trial was not rewarded. **(c)** Lever position vs. time traces (n = 50 trials) for a representative WT mouse on the seventh day of testing. Histogram indicates the amplitude of displayed trials and grey-shaded area indicates the rewarded pull amplitude range. **(d)** WT mice (*n* = 13) increased their proportion of rewarded trials (*p* < 0.0001) and decreased their proportion of overshot trials (*p* < 0.0001) and trials held for longer than the trial time limit (*p* < 0.0001) over one week of testing. The proportion of undershot trials remained relatively stable (*p* = 0.32). **(e)** The proportion of rewarded trials did not increase significantly past day 7 of testing in a group of WT mice (*n* = 7) assessed on the task for three weeks. **(f)** Kernel density estimate (KDE) plot of pull amplitude across trials for all WT mice (*n* = 13) on the first and seventh day of testing. **(g)** The average pull amplitude for WT mice (*n* = 13) over one week of testing did not change significantly (*p* = 0.12). The grey shaded region indicates the rewarded pull amplitude range. **(h)** The Fano factor 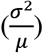 of pull amplitude calculated for each day decreased significantly over one week of testing (*p* < 0.0001) in WT mice (*n* = 13). **(i)** The Fano factor of pull amplitude for each mouse on each day was correlated with daily reward rate for that mouse (*n* = 91 mouse-days). All data are presented as mean ± SEM.

The lever was coupled to a DC micro-motor, which outputted a fixed torque in order to hold the lever in its ‘start’ position (towards the spout). The torque was set to one of two levels: a ‘low-force’ condition when a mouse was performing a trial, and a ‘high-force’ condition at all other times (e.g. during the time-out between trials). The torque applied during the low-force condition was very low (~15 mN) – the minimum required to overcome the friction of the rotor and return the lever back to the start position after it was pulled backwards. Two micro-motors were used for the described experiments – a 15mm model (Faulhaber 1524T012SR) and a larger 22mm model (Faulhaber 2224U012SR). As the 22mm motor was mounted slightly further away from the testing module, the absolute size of each degree of rotation at the end of the lever was larger with this motor as compared to the 15mm model. In order to compensate for this difference and maintain the same arc length of lever movement between the two setups, the size of the lever range was decreased to 24° from 30° for all cages tested with the large motor (all sub-ranges were scaled proportionately as well). During analysis, all lever position data for animals tested with the large motor were scaled by a factor of 1.25× in order to normalize the data to the 30° range used with the small motor. No differences in pull kinematics or learning were seen between cohorts tested in cages with the two different motors. The motor was coupled to a high resolution (4096 lines per revolution) incremental encoder (Faulhaber IEH2-4096) to allow for accurate measurement and recording of all lever movements. Lever positions during each trial were collected at 200 Hz. In addition, a camera (Waveshare 10299) was mounted below the chamber and recorded a bottom-up video of each trial through the glass floor. A piezo buzzer delivered auditory stimuli to indicate trial initiation, reward and failure. All components were connected to a breakout board and controlled by custom software running on a Raspberry Pi 3B micro-computer. This software was written in Python and is available online (https://github.com/cameron-woodard/PiPaw).

#### Testing methodology (amplitude task)

Small groups (<5) of littermate mice were introduced to the PiPaw cage and allowed to explore and discover the attached testing module. *Ad libitum* water bottle access was removed, so that water could only be retrieved by interacting with the behavioural task. In order to initiate a trial, the mouse had to nose-poke at the port and remain in position for a short waiting period (1 s). Following this, the motor switched to the low-force condition and a short tone was played. The mouse could then initiate a trial by pulling the lever backwards out of the ‘threshold’ position range (0° to 3). The trial was ended either when the lever was returned to the start position, or when the trial time limit of 2 s was reached, after which there was a 5 s timeout before the next trail could begin. If the reward requirements were met, a 20 μL water drop was delivered by the spout.

Testing was split into three phases which were completed sequentially, each animal advancing at their own pace (Fig. 1b). In the first phase (Training 1) mice acquired the operant response of nose-poking and simultaneously pulling the lever with their right forelimb. Mice received a water drop reward on a fixed-interval 15-minute (FI-15) schedule for nose-poking, and could receive additional rewards by pulling the lever past the threshold range and returning it to the start position (either intentionally or by letting go of it) within the 2 s trial time limit. A trial could only be failed by holding the lever too long, encouraging mice to perform short movements. Once a mouse reached 100 rewarded trials in this phase, they were advanced to Training 2. In this phase, mice no longer received rewards simply for nose poking, and had to pull the lever past 8° in order to be rewarded. After 100 rewarded trials in this phase, mice were moved on to the main Testing phase. In the Testing phase, trials were rewarded if the amplitude of the pull (i.e. the maximum displacement of the lever from the ‘start’ position) was between 15° and 27° of the full 30° lever position range. As in the Training phases, if the lever was held for longer than 2 s, the trial was also not rewarded. Mice were assessed in this main Testing phase for between one to four weeks. In all phases, when the lever entered the rewarded position range (i.e. when it passed 3° in Training 1, 8° in Training 2 or 15° in Testing), a short high tone was played. In the Testing phase, if the mouse pulled the lever back past the far end of the rewarded position range (27°), a short low tone was played. These tones served to reinforce the location of the rewarded range within the full lever position range.

#### Testing methodology (velocity task)

The procedure for introducing mice to the testing system and the general structure of trials was the same as with the amplitude task. The first training phase (Training 1) was also the same as the amplitude task, with all pulls rewarded as long as the lever was returned to the start position within the 2 s trial time limit. Once a mouse reached 100 rewarded trials in Training 1, they were advanced to Training 2, in which trials were rewarded only if the amplitude of the pull was greater than 6°. After a further 100 rewarded trials, each mouse was advanced to the main Testing phase. In the Testing phase, trials were rewarded if the amplitude of the pull was greater than 6° and the average velocity of the pull to the maximum lever position was between 50 and 100 degrees per second. If the pull velocity was less than 50 °/s or greater than 100 °/s, no reward was delivered. As in the Training phases, if the lever was held for longer than 2 s, the trial was also not rewarded. Mice were assessed in this main Testing phase for three weeks.

#### Data analysis

All data were automatically recorded to text files by the PiPaw software and were extracted and analyzed using custom scripts written in Python. Prior to analysis, lever position data were ‘cleaned’ in order to remove trials with abnormal timestamps or lever position readings, as well as trials with 2 or fewer total position readings. This cleaning resulted in the removal of only a very small number of trials (~0.1%). To perform daily analysis of task performance, trials were grouped into 24-hour bins from the time that the animal was switched to the main Testing phase. These bins were used to determine the mean and variance of kinematic measures, and the Fano factor was calculated as the variance divided by the mean for each daily bin. To define bouts of trials, the average trial performance rate was calculated for a 3-minute sliding window across the full Testing phase for each animal. When this rate went above 1.333 trials/min (corresponding to >4 trials in the 3-minute window), all trials in the window were grouped into a bout and the bout continued for as long as the trial rate stayed above this value. Although the design of the system encouraged mice to pull the lever with their right forelimb, a small but significant proportion of animals (~18% of mice) performed trials with either their left forelimb or both forelimbs. As the use of a different pull strategy on some trials could be a confounding factor for kinematic analysis, these mice were excluded from analysis. In order to identify these animals, 50 trial videos were randomly selected for each animal and manually scored for pull strategy by a blinded observer. Mice that had greater than 3/50 videos scored as left or both forelimbs were excluded from analysis. In addition, a small number of animals (*n* = 3) were excluded from analysis as they were assessed for less than seven days in the main Testing phase.

### Accelerating rotarod test

In the rotarod test, a mouse is placed on a rotating rod and must balance themselves in order to prevent from falling. On each trial, the rotarod (Ugo Basile, Italy) accelerated from 5 to 40 RPM over the course of 300 s, and the latency for each mouse to fall from the rotarod was noted. If the mouse performed a complete rotation holding onto the rod, this was also treated as a fall and the trial was ended. If the mouse reached the maximum allowed time, the trial was ended and scored as 300 s. The rotarod was wiped with ethanol between each mouse. Testing was performed at the same time on each day during the light phase of the light/dark cycle. Each mouse performed 3 trials per day on each of 4 consecutive days, with a one-hour inter-trial interval. Latency to fall off the rotarod on the three trials each day was averaged for each mouse to obtain a daily average.

### Electrophysiology

Electrophysiology experiments were performed on striatal slices collected from 10- to 11-month-old mice that either had not undergone PiPaw testing or who had been trained on the PiPaw amplitude task. Mice in the trained group had been tested for 3-4 weeks and were removed from the PiPaw cage immediately prior to terminal experiments.

#### Slice preparation

Animals were anesthetized with isoflurane and decapitated. The brain was rapidly removed and bisected along the midline, separating the two hemispheres. Acute left- and right-hemisphere sagittal slices (250-300 μm) containing the dorsal striatum were cut using a vibratome (Leica VT1000) in ice-cold artificial cerebrospinal fluid (aCSF), before being transferred to a holding chamber containing aCSF at 37° for 30 minutes. Slices were then maintained in aCSF at room temperature for at least 30 minutes for whole-cell experiments, or 1 hour for extracellular experiments. All aCSF contained the following (in mM): 125 NaCl, 2.5 KCl, 25 NaHCO3, 1.25 NaH2PO4 and 10 glucose. In addition, aCSF used for cutting slices contained 0.5mM CaCl2 and 2.5mM MgCl2, while all other aCSF contained 2mM CaCl2 and 1mM MgCl2. The pH of aCSF was 7.3-7.4 and osmolarity was 310 (±3) mOsm/L. aCSF was continuously oxygenated with carbogen (95% O2/5% CO2) during slicing, recovery and all experiments. Once transferred to the recording chamber, slices were continuously superfused with room temperature aCSF containing picrotoxin (50μM; Tocris Bioscience) to block GABA_A_ receptors and minimize inhibitory responses. Slices were allowed to equilibrate in the recording chamber for at least 20 minutes before the start of recording.

#### Whole-cell voltage-clamp

Intracellular recordings were made using a whole-cell patch clamp technique and were acquired with an Axopatch-700A amplifier and pClamp 11 software, digitized at 20kHz and filtered at 1 kHz. Pipettes (3-5Ω) were pulled from borosilicate glass capillaries using a micropipette puller (Narishige International). The intracellular solution was cesium-based and contained the following in mM: 130 cesium methanesulfonate, 5 CsCl, 4 NaCl, 1 MgCl2, 5 EGTA, 10 HEPES, 5 QX-314 chloride, 5 MgATP, 0.5 MgGTP and 10 sodium phosphocreatine. The pH of intracellular solution was 7.2-7.3 and the osmolarity was 290 (±3) mOsm/L. Cells were rejected and not recorded if series resistance was >17 MΩ. To record spontaneous excitatory post-synaptic currents (sEPSCs), cells were voltage-clamped at −70 mV. To record the paired-pulse ratio (PPR), a glass micropipette electrode filled with aCSF was positioned ~200 μm dorsal to the recording site. The cell was voltage clamped at −70 mV with a 50 ms step to −80 mV every 30 s, and activity was elicited by injecting current through the stimulating electrode. Two pulses (100 μs duration) were administered with an inter-pulse interval of either 50, 100, 150, 200 or 250 ms. Three runs were performed at each interval length and averaged, and the PPR at each interval was calculated as the ratio of the average response amplitude of the second pulse to the average response amplitude of the first pulse. Analysis of electrophysiology data was performed using Clampfit 10.7 (Molecular Devices).

### Experimental design and statistical analysis

The experimenter was blinded to genotype during all experiments. Statistical testing was performed using Prism 8 (GraphPad Software) and Python. Data are expressed as mean ± SEM unless otherwise specified. Alpha level for all tests was *p* = 0.05. For electrophysiology experiments, *n* = the number of neurons and the number of animals is given in brackets. For all other experiments, *n* = the number of animals. Repeated measures data with group comparisons were analyzed with repeated measures two-way ANOVA to assess overall main and interaction effects, followed by Sidak’s multiple comparisons test to compare between groups at each time-point. Repeated measures data for a single group were analyzed using repeated measures one-way ANOVA followed by Tukey’s test to compare timepoints. To compare non-repeated measures data across three or more groups, one-way ANOVA was performed followed by Tukey’s test to compare pairs of groups. To compare two groups on a single measure, unpaired two-tailed t-tests were used when groups were normally distributed and had equal variances. If groups were found to have unequal variances using the F-test of equality of variances, Welch’s t-test was used instead. If one or both groups were found to have a non-normal distribution using the D’Agostino & Pearson test, the Mann-Whitney test, a non-parametric alternative, was used. To compare paired data, paired two-tailed t-tests were used. Pearson correlation coefficients were calculated to measure the linear correlation between two measures.

## Results

### Mice improve their performance of a skilled forelimb task by reducing the inter-trial variability of reward-related kinematic parameters

Using the PiPaw home-cage task (Fig. 1a), we first assessed forelimb motor learning in a cohort of wildtype mice (*n* = 13). Initially, mice were rewarded with water drops on a fixed-interval 15-minute schedule (FI-15) for nose-poking at the port and could obtain additional drops by simultaneously pulling the lever with their forelimb (continuously reinforced). Pulls of any amplitude were rewarded during the first training phase, while in the second training phase, only pulls with an amplitude of >8° resulted in water drop delivery (Fig. 1b). Once a mouse had acquired this operant response, they were automatically advanced to the main testing phase, in which trials were rewarded only if the amplitude of the pull was between 15° and 27° (Fig. 1b-c). In this phase, the proportion of rewarded trials was initially quite low (19.0% ± 3.7% on day 1), but increased significantly over the course of one week, reaching 50.0% ± 2.6% by day 7 (*F*_6, 72_ = 25.6, *p* < 0.0001, ANOVA) (Fig. 1d). This improvement in task performance was accompanied by a concomitant decrease in the number of ‘overshot’ trials (amplitude of >27°) (*F*_6, 72_ = 7.20, *p* < 0.0001, ANOVA) and trials held for longer than the time limit of 2 seconds (*F*_6, 72_ = 15.1, *p* < 0.0001, ANOVA) (Fig. 1d). In a subset of mice tested on the task for an additional two weeks (*n* = 7), the proportion of rewarded trials continued to increase (64.5% ± 3.8% on day 21). However, reward rate was not significantly different on day 21 as compared to day 7 (*p* = 0.06, Tukey), indicating that performance gains occurred primarily in the first week of testing (Fig. 1e).

As mice improved their performance of the task over one week, the distribution of pull amplitude across trials changed substantially (Fig. 1f). Interestingly, the mean amplitude of pulls was relatively stable over seven days (*F*_6, 72_ = 1.74, *p* = 0.12, ANOVA) and was within the rewarded range even on the first day of testing, suggesting that performance improvements were not primarily driven by an increase or decrease in average pull amplitude (Fig. 1g). However, the inter-trial variability of pull amplitude (quantified as the Fano factor 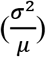 for each 24-hour bin) decreased by almost 40% over the course of one week (*F*_6, 72_ = 11.4, *p* < 0.0001, ANOVA) (Fig. 1h). In addition, the Fano factor of pull amplitude had a strong negative correlation with daily reward rate for each animal (*r* = −0.73, *p* < 0.0001, Pearson; *n* = 91), suggesting that the observed reduction in movement variability was directly related to improved performance of the task (Fig. 1i). We next investigated whether this reduction in kinematic variability was specific to reward-related parameters (i.e. pull amplitude) or instead reflected a general increase in movement stereotypy. Mice also showed a decrease in the Fano factor of peak pull velocity over one week of testing (*F*_6, 72_ = 13.2, *p* < 0.0001, ANOVA); however, the Fano factor of peak acceleration did not change (*F*_6, 72_ = 0.73, *p* = 0.63, ANOVA) (data not shown). This suggests that kinematic variability is not universally reduced with training and is instead regulated in a more specific manner.

To further investigate the relationship between refinement of kinematic variability and reward, we tested an additional cohort of mice (*n* = 16) on a task where trials were rewarded if the lever was pulled with a specific average velocity (50-100 °/s) rather than with a specific amplitude (Fig. 2a). As with the amplitude task, mice performing the velocity task improved their performance over time (*F*_6, 90_ = 6.28, *p* < 0.0001, ANOVA), although the proportion of rewarded trials increased comparatively gradually (Fig. 2b). Average reward rate did not increase substantially after day 7 (*p* = 0.67, Tukey), plateauing at an average of 39.1% ± 3.3% after three weeks of testing (Fig. 2c). Interestingly, mice did not decrease the variability of pull amplitude over time in this task; rather, the Fano factor of this parameter increased on average across the first week of testing (*F*_6, 90_ = 2.28, *p* = 0.04, ANOVA). In contrast, the inter-trial variability of peak pull velocity decreased significantly over one week (*F*_6, 90_ = 3.32, *p* = 0.005, ANOVA). This suggests that mice can independently regulate the variability of kinematic parameters while interacting with the task and prioritize reducing variability in parameters most relevant to reward.

**Figure 2:**
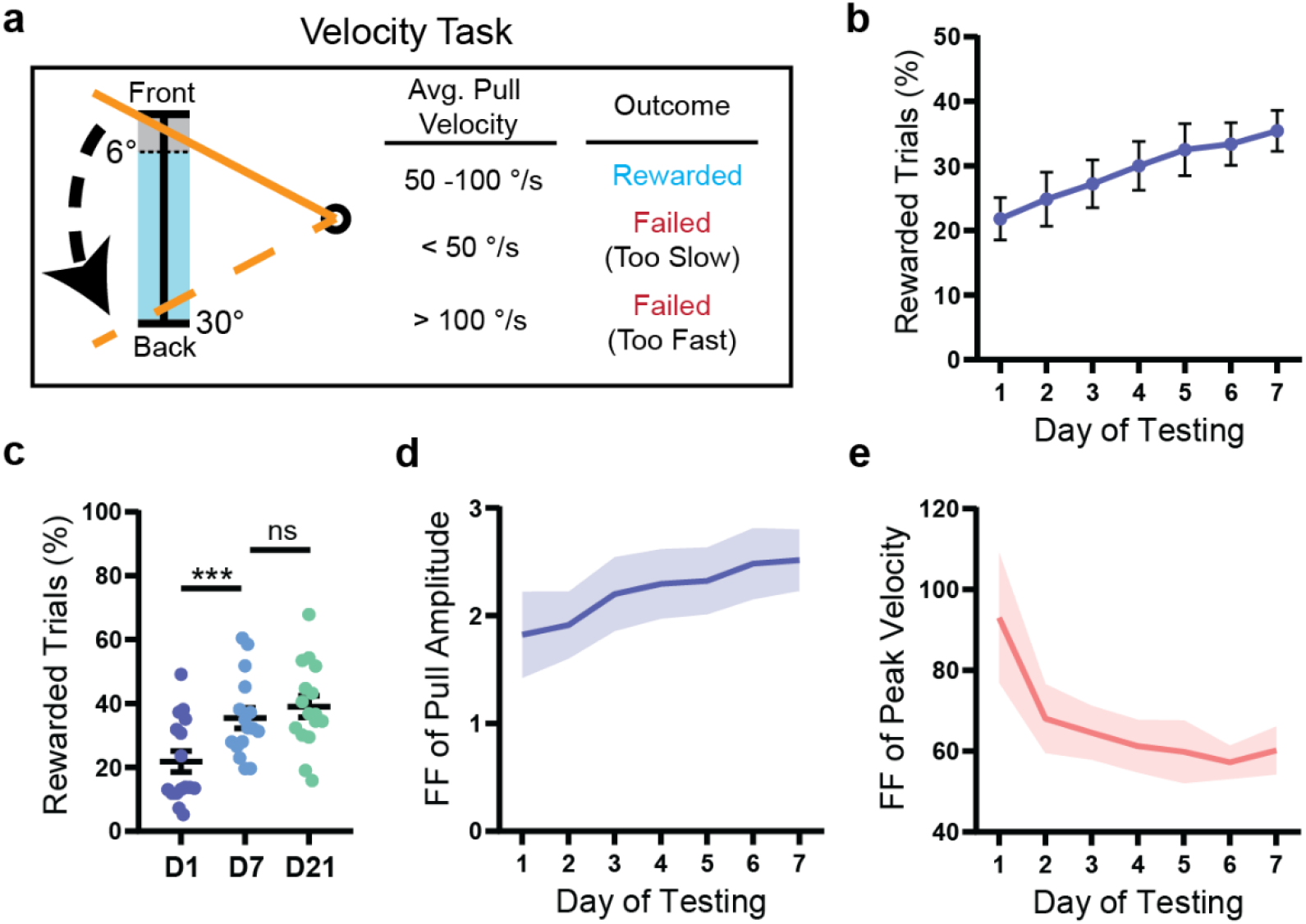
Mice preferentially reduce the inter-trial variability of reward-related kinematic parameters. **(a)** In the velocity task, a trial was rewarded if the average velocity of the lever (to its maximum position) was between 50 and 100 degrees per second. Mice initiated a trial by pulling the lever past the threshold range (0-6°, shown in grey) and were required to return the lever to the start position within 2 s. **(b)** WT mice (*n* = 16) improved their performance of the task over one week (*p* < 0.0001). **(c)** The proportion of rewarded trials did not increase significantly past day 7 of testing in WT mice (*n* = 16). **(d)** The Fano factor of pull amplitude calculated for each day of testing increased significantly over one week (*p* = 0.04) in WT mice (*n* = 16). **(e)** The Fano factor of peak velocity calculated for each day of testing decreased significantly over one week (*p* = 0.005) in WT mice (*n* = 16). All data are from mice tested on the velocity task and are presented as mean ± SEM.

### Mice organize their activity into short bouts of high task engagement

Mice assessed on the PiPaw task performed an average of 251.4 ± 11.6 trials per day during the main testing phase. A circadian pattern of task engagement was observed, with activity increasing in the late afternoon and peaking during the first three hours of the dark phase between 7 and 9 PM before decreasing (Fig. 3a). In addition, when mice interacted with the task, they tended to cluster their trials into short (<10 minutes) ‘bouts’ of high task engagement, rather than distributing them more evenly over time. To identify and segregate these trial bouts, we determined the average trial rate in a three-minute sliding window for each mouse and grouped trials when the rate exceeded a specified value (>1.333 trials/minute). Using this method, we found that 92.3% ± 0.9% of each mouse’s trials occurred in these periods of high task engagement, with the remainder showing a sparser distribution (Fig. 3b). Mice performed an average of 18.7 ± 0.7 bouts per day, with each bout containing an average of 12.5 ± 0.5 trials. Interestingly, the reward rate in trials occurring within bouts was found to be significantly higher than the reward rate in trials occurring outside of bouts (*t*_12_ = 6.16, p < 0.0001, paired t-test; *n* = 13) (Fig. 3c). One possible explanation for this is that clustering trials into bouts of activity facilitates short-term improvements in task performance. In support of this, we found that the average reward rate (assessed over a 5-trial window) increased over the course of each bout, with trials later in the bout having a higher reward rate than those at the beginning (*F*_5, 60_ = 10.89, *p* < 0.0001, ANOVA; *n* = 13) (Fig. 3d). Furthermore, this increase in reward rate was paralleled by a within-session decrease in the inter-trial variability of pull amplitude (5-trial window) (*F*_5, 60_ = 6.67, *p* < 0.0001, ANOVA; *n* = 13) (Fig. 3e). This indicates that mice reduce the variability of their movements on both short and long timescales in order to improve their performance of the task.

**Figure 3:**
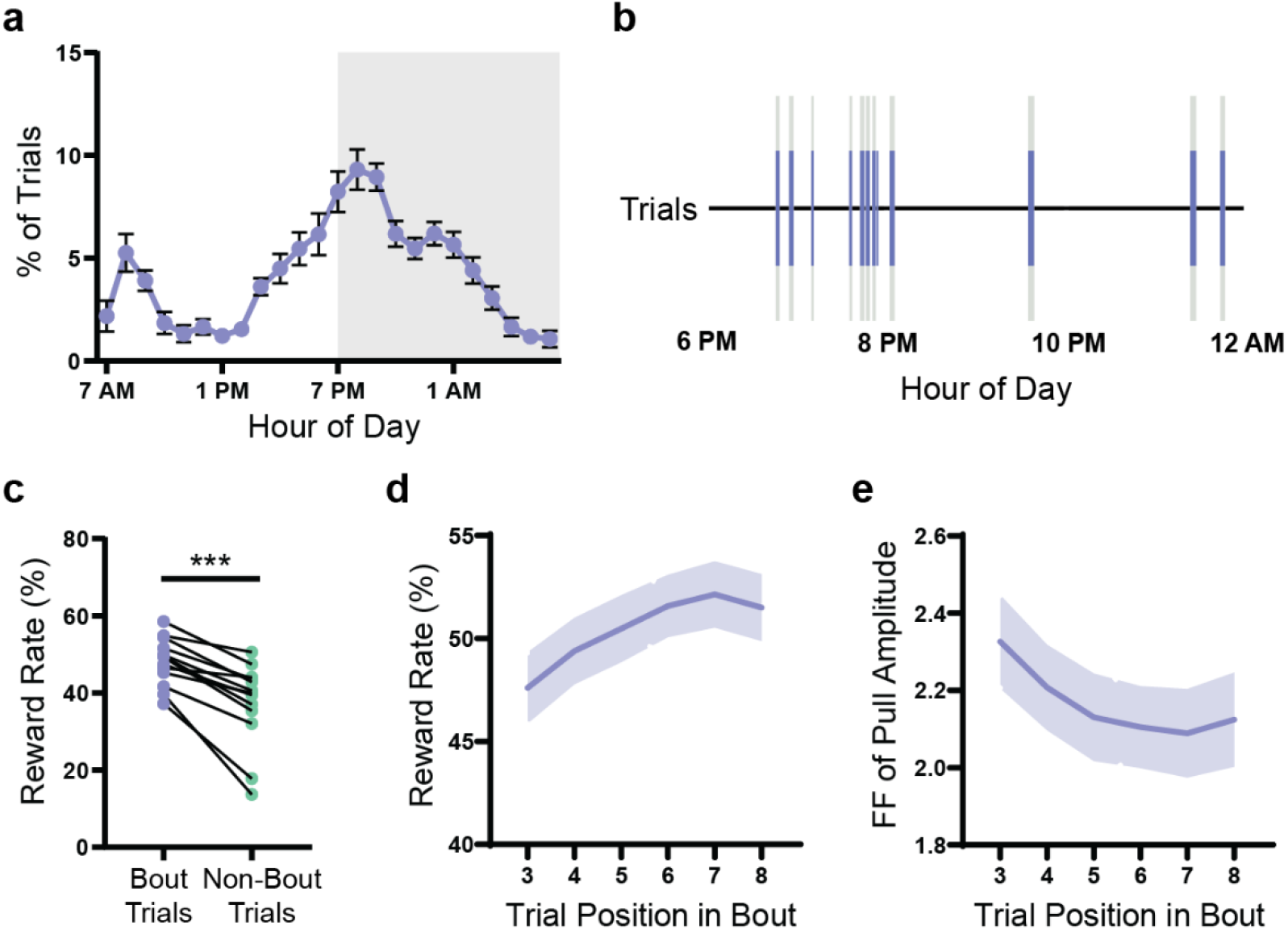
Mice organize their activity into short bouts of high task-engagement. **(a)** Average circadian distribution of trials (hourly bins) in WT mice (*n* = 13). Grey-shaded region indicates the dark phase of the light cycle (7 PM – 7 AM). **(b)** Trials (blue lines) for a representative mouse are shown over an 8-hour period. The large majority of trials occurred in bouts (grey-shaded regions) of high task engagement (>4 trials within a 3-minute window). **(c)** Reward rate was higher for trials occurring within bouts as compared to trials occurring outside of bouts (*p* < 0.0001). Lines indicate paired values for each WT mouse (*n* = 13). **(d)** Reward rate increased over the course of a bout (*p* < 0.0001) in WT mice (*n* = 13). Reward rate was calculated for a 5-trial window centered on the indicated trial position (e.g. trials 1-5 for trial 3). **(e)** The Fano factor of pull amplitude decreased over the course of a bout (*p* < 0.0001) in WT mice (*n* = 13) (amplitude task). Fano factor was calculated for a 5-trial window centered on the indicated trial position. All data are from mice tested on the amplitude task and are presented as mean ± SEM.

### Q175-FDN HD mice have impaired forelimb motor learning that is not explained by differences in mean pull kinematics

We next used the PiPaw task to assess motor learning in two mouse models of Huntington disease. The transgenic YAC128 model and the knock-in Q175-FDN model both have a slowly progressing phenotype, with motor coordination impairments first emerging around 6-months-old in YAC128 mice (Slow et al., 2003) and 8-months-old in Q175-FDN mice (Southwell et al., 2016). At 10- to 11-months-old, YAC128 (*n* = 11) and Q175-FDN mice (*n* = 10) had significantly impaired performance on the accelerating rotarod task over four days of testing as compared to WT mice (*n* = 11) (Day: *F*_3, 87_ = 24.8, *p* < 0.0001; Genotype: *F*_2, 29_ = 9.42, *p* = 0.0007; Interaction: *F*_6, 87_ = 2.59, *p* = 0.02; ANOVA) (Fig. 4a), and the degree of motor impairments was similar between the two HD models (YAC128 vs. Q175-FDN: *p* = 0.60, Tukey). Surprisingly, when YAC128 mice (*n* = 15) were assessed in the PiPaw task, they had normal motor learning and improved at a similar rate to WT controls (*n* = 28) over one week of testing (Day: *F*_6, 246_ = 53.4, *p* < 0.0001; Genotype: *F*_1, 41_ = 0.003, *p* = 0.96; Interaction: *F*_6, 246_ = 0.21, *p* = 0.97; ANOVA) (Fig. 4b). In contrast, Q175-FDN mice (*n* = 14) were significantly impaired at learning this forelimb task (Day: *F*_6, 240_ = 30.4, *p* < 0.0001; Genotype: *F*_1, 40_ = 18.1, *p* = 0.0001; Interaction: *F*_6, 240_ = 7.42, *p* < 0.0001; ANOVA) (Fig. 4b).

**Figure 4:**
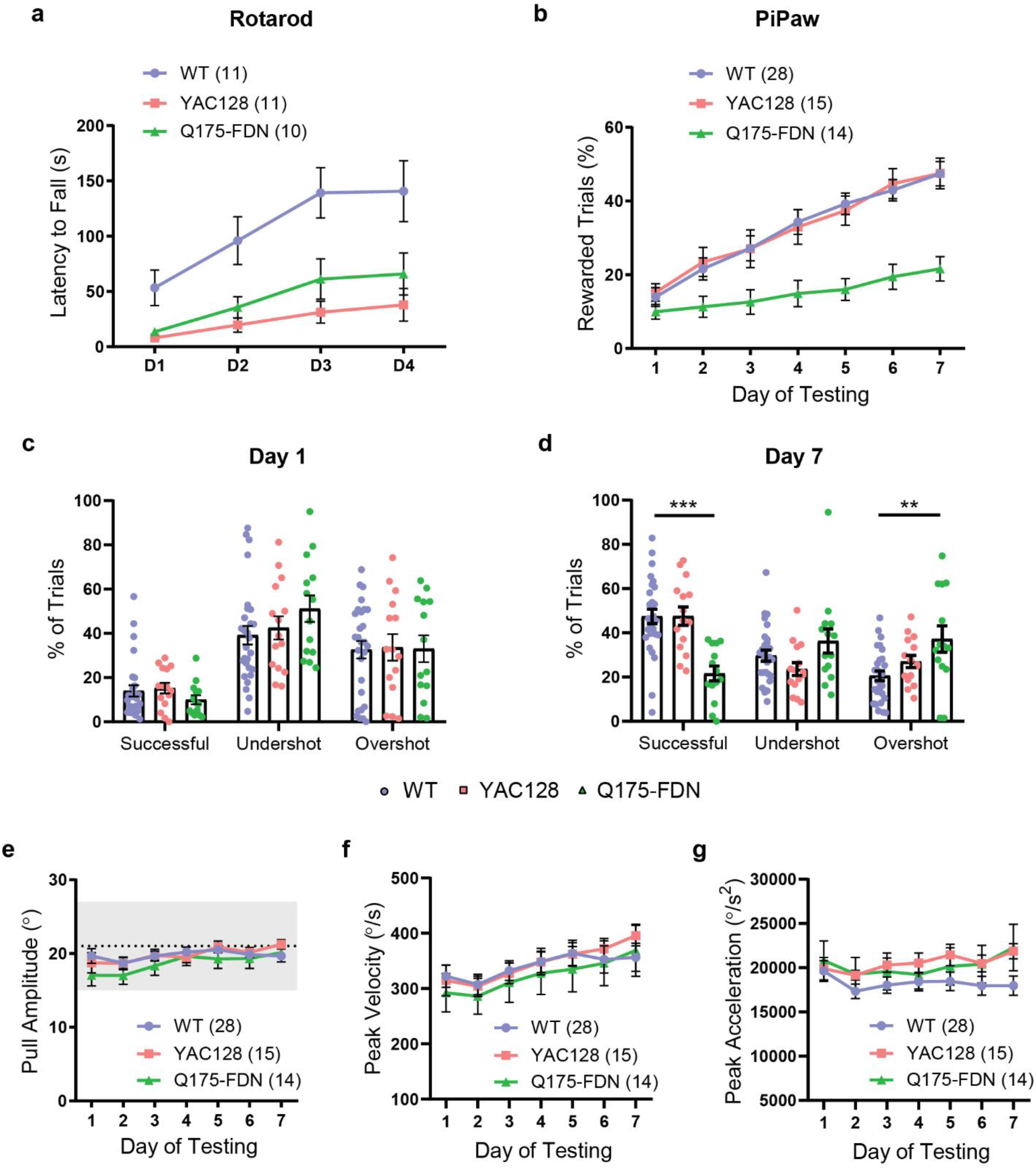
Q175-FDN HD mice have impaired forelimb motor learning not explained by differences in mean pull kinematics. **(a)** YAC128 and Q175-FDN HD mice (10-months-old) have impaired performance of the accelerating rotarod task over four days of testing (WT vs. YAC128: *p* = 0.0007; WT vs. Q175-FDN: *p* = 0.012). **(b)** Q175-FDN, but not YAC128, mice (10-months-old) have impaired learning of the PiPaw amplitude task (WT vs. YAC128: *p* = 0.99; WT vs. Q175-FDN: *p* < 0.0001). **(c)** WT, YAC128 and Q175-FDN mice have a similar proportion of rewarded, undershot and overshot trials on the first day of testing. **(d)** By day 7 of testing, Q175-FDN mice have fewer rewarded trials (*p* < 0.0001) and more overshot trials (*p* < 0.003) as compared to WT mice. **(e)** Average pull amplitude is not different between genotypes across one week of testing. **(f)** Average peak pull velocity is not different between genotypes across one week of testing. **(g)**Average peak pull acceleration is not different between genotypes across one week of testing. All PiPaw data are from mice tested on the amplitude task and are presented as mean ± SEM.

The impairment observed in Q175-FDN mice was not related to baseline differences in task performance, as the proportion of rewarded, undershot and overshot trials was similar between the three genotypes on the first day of testing (Trial Outcome: *F*_2, 162_ = 35.9, *p* < 0.0001; Genotype: *F*_2, 162_ = 0.35, *p* = 0.71; Interaction: *F*_4, 162_ = 0.9235, *p* = 0.45; ANOVA) (Fig. 4c). By the seventh day of testing, however, a significant interaction was seen between genotype and trial outcome (Trial Outcome: *F*_2, 162_ = 7.3, *p* = 0.0009; Genotype: *F*_2, 162_ = 0.06, *p* = 0.94; Interaction: *F*_4, 162_ = 11.64, *p* < 0.0001; ANOVA), and Q175-FDN mice had a lower proportion of rewarded trials and a higher proportion of overshot trials as compared to WT controls (Fig. 4d). Impaired learning of the task in Q175-FDN mice was not related to differences in the average amplitude of pulls, as this was similar between genotypes on all days of testing (Day: *F*_6, 324_ = 5.97, *p* < 0.0001; Genotype: *F*_2, 54_ = 0.69, *p* = 0.50; Interaction: *F*_12, 324_ = 1.05, *p* = 0.40; ANOVA) (Fig. 4e). Similarly, no differences were seen between WT, YAC128 and Q175-FDN mice in either the average peak velocity (Day: *F*_6, 324_ = 9.21, *p* < 0.0001; Genotype: *F*_2, 54_ = 0.20, *p* = 0.82; Interaction: *F*_12, 324_ = 0.46, *p* = 0.94; ANOVA) (Fig. 4f) or average peak acceleration of pulls across testing (Day: *F*_6, 324_ = 2.49, *p* = 0.02; Genotype: *F*_2, 54_ = 1.27, *p* = 0.29; Interaction: *F*_12, 324_ = 1.11, *p* = 0.35; ANOVA) (Fig. 4g). This suggests that impaired performance of the task in Q175-FDN mice was not caused by overt differences in mean movement kinematics.

### Q175-FDN HD mice are unable to decrease movement variability in order to improve task performance

To further investigate impaired motor learning in Q175-FDN mice, we next examined the distribution of pull amplitude across trials. Similarly to the cohort of 2-month-old WT mice, WT mice from the 10-month-old cohort shifted the distribution of pull amplitude across days, such that the probability of a trial having either a very high or a very low amplitude decreased, and the probability of a trial having an amplitude in the central rewarded range increased by the seventh day of testing (Fig 5a-b). In Q175-FDN mice, however, this shift was subtle, and was characterized by an increased probability of higher pull amplitude generally, rather than within the goal range specifically, by the seventh day of testing (Fig. 5c-d). The Fano factor of pull amplitude (calculated for each day of testing) was significantly higher in Q175-FDN as compared to WT mice (Fig. 5e). However, this measure decreased over time in both groups (Day: *F*_6, 240_ = 17.6, *p* < 0.0001; Genotype: *F*_1, 40_ = 4.77, *p* = 0.04; Interaction: *F*_6, 240_ = 0.31, *p* = 0.93; ANOVA), suggesting that Q175-FDN mice were able to reduce the variability of their movements to a certain extent (Fig. 5e).

**Figure 5:**
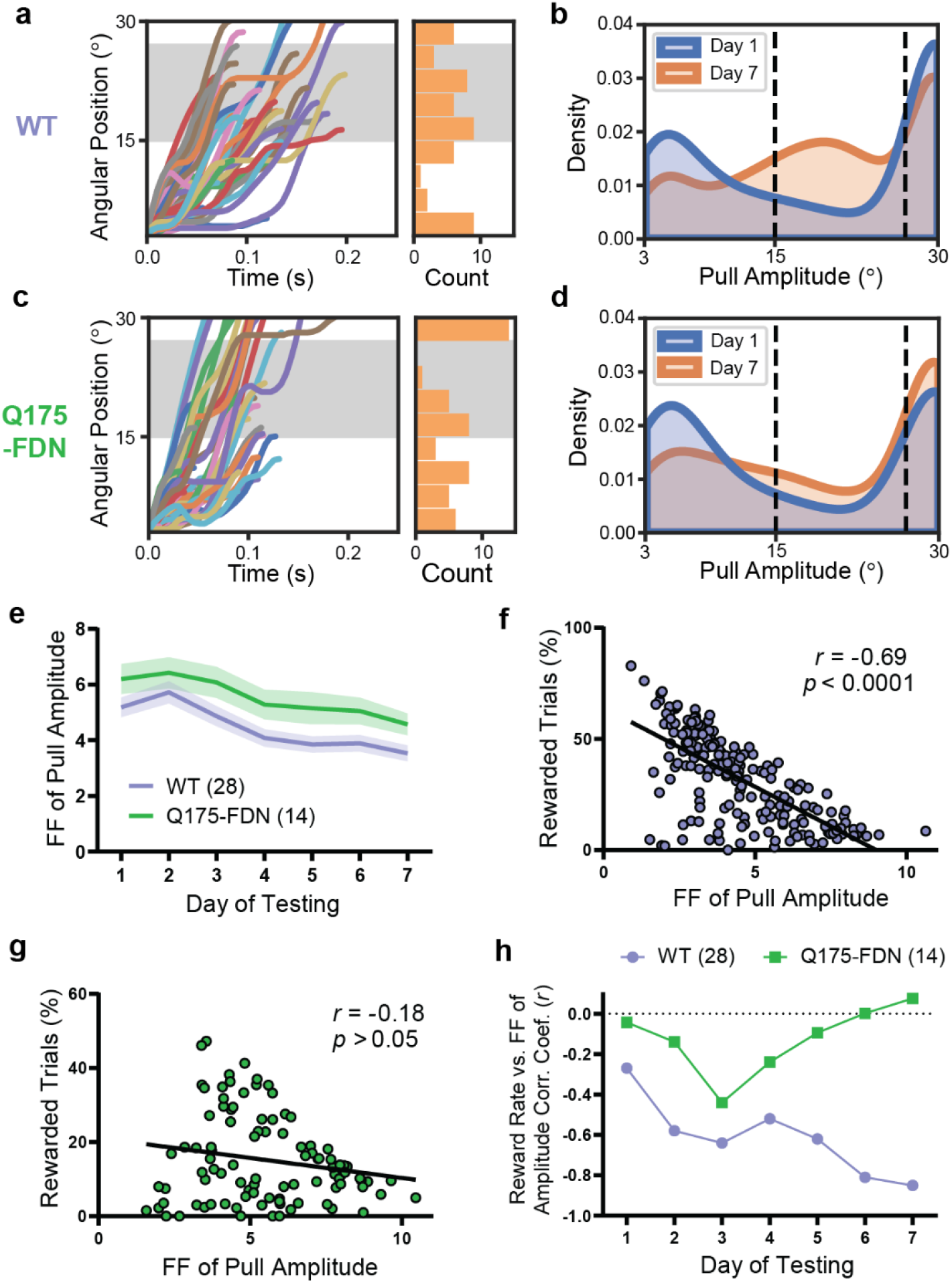
Q175-FDN mice are unable to modulate the inter-trial variability of pull amplitude in order to improve task performance. **(a)** Lever position vs. time traces (*n* = 50 trials) for a representative WT mouse on the seventh day of testing. Histogram indicates the amplitude of displayed trials and grey-shaded area indicates the rewarded pull amplitude range. **(b)** Kernel density estimate (KDE) plot of pull amplitude across trials for all WT mice (*n* = 28) on the first and seventh day of testing. **(c)** Lever position vs. time traces (*n* = 50 trials) for a representative Q175-FDN mouse on the seventh day of testing. **(d)** KDE plot of pull amplitude across trials for all Q175-FDN mice (*n* = 14) on the first and seventh day of testing. **(e)** The Fano factor of pull amplitude calculated for each day was significantly higher in Q175-FDN as compared to WT mice (*p* = 0.035), although it decreased over time in both genotypes (*p* < 0.0001). **(f)** The Fano factor of pull amplitude for each WT mouse on each day was correlated with daily reward rate for that mouse (*n* = 196 mouse-days). **(g)** Fano factor of pull amplitude was not correlated with daily reward rate in Q175-FDN mice (*n* = 98 mouse-days). **(h)** Correlation coefficient of Fano factor of pull amplitude and reward rate for WT and Q175-FDN mice across seven days of testing. The correlation is significant for WT mice from day 2 onwards but is not significant in Q175-FDN mice on any day. All data are from mice tested on the amplitude task and are presented as mean ± SEM.

To investigate whether inter-trial variability of pull amplitude was related to task performance on an individual level, we next measured the linear correlation between reward rate and variability of pull amplitude across animals. Overall, the Fano factor of pull amplitude was inversely correlated with daily reward rate only in WT mice (*r* = −0.69, *p* < 0.0001, Pearson; *n = n =* 196) (Fig. 5f) and not in Q175-FDN animals (*r* = −0.18, *p* = 0.07, Pearson; *n* = 98) (Fig. 5g). Interestingly, calculation of the correlation coefficient of amplitude variability and reward rate showed that these parameters were not correlated on the first day of testing in either genotype (WT: *r* = −0.28, *p* = 0.14; Q175-FDN: *r* = −0.04, *p* = 0.88; Pearson; *n* = 28 WT, 14 Q175-FDN) (Fig. 5h). This indicates that mice who performed pulls with a more consistent amplitude initially did not necessarily receive more rewards (and indeed may have received fewer rewards if their average amplitude was too high or too low). However, amplitude variability and reward rate became progressively more negatively correlated in the WT group and were highly correlated by the seventh day of testing (*r* = −0.88, *p* < 0.0001, Pearson; *n* = 28) (Fig. 5h). This pattern was not seen in Q175-FDN mice, and no significant correlation between these two measures was observed on any day of testing in this group (Fig. 5h). This suggests that the decrease in inter-trial variability of pull amplitude observed on a group level in Q175-FDN mice was not related to individual improvements in task performance.

We next examined whether patterns of task engagement and circadian activity levels were different in Q175-FDN as compared to WT animals. Consistent with reports in other HD mouse models (Morton et al., 2005; Kudo et al., 2011; Loh et al., 2013; Woodard et al., 2017), Q175-FDN mice had a significantly altered circadian distribution of their trials as compared to WT littermates (Hour: *F*_23, 920_ = 44.56, *p* < 0.0001; Genotype: *F*_1, 40_ = 0.078, *p* = 0.78; Interaction: *F*_23, 920_ = 6.53, *p* < 0.0001; ANOVA), with a greater proportion of trials occurring in the second half of the dark phase (1-7 AM) (Fig. 6a). Q175-FDN mice also performed a greater number of trials each day on average (*U* = 121, *p* = 0.046, Mann-Whitney), likely as a result of this group requiring more trials in order to maintain adequate daily water intake (due to their lower reward rate) (Fig. 6b). This increased level of task engagement did not manifest as a greater number of trials in each bout (*U =* 190, *p* = 0.88, Mann-Whitney) (data not shown), but rather as an increased number of trial bouts per day (*t*_17.4_ = 2.99, *p* = 0.008, t-test) (Fig. 6c).

**Figure 6:**
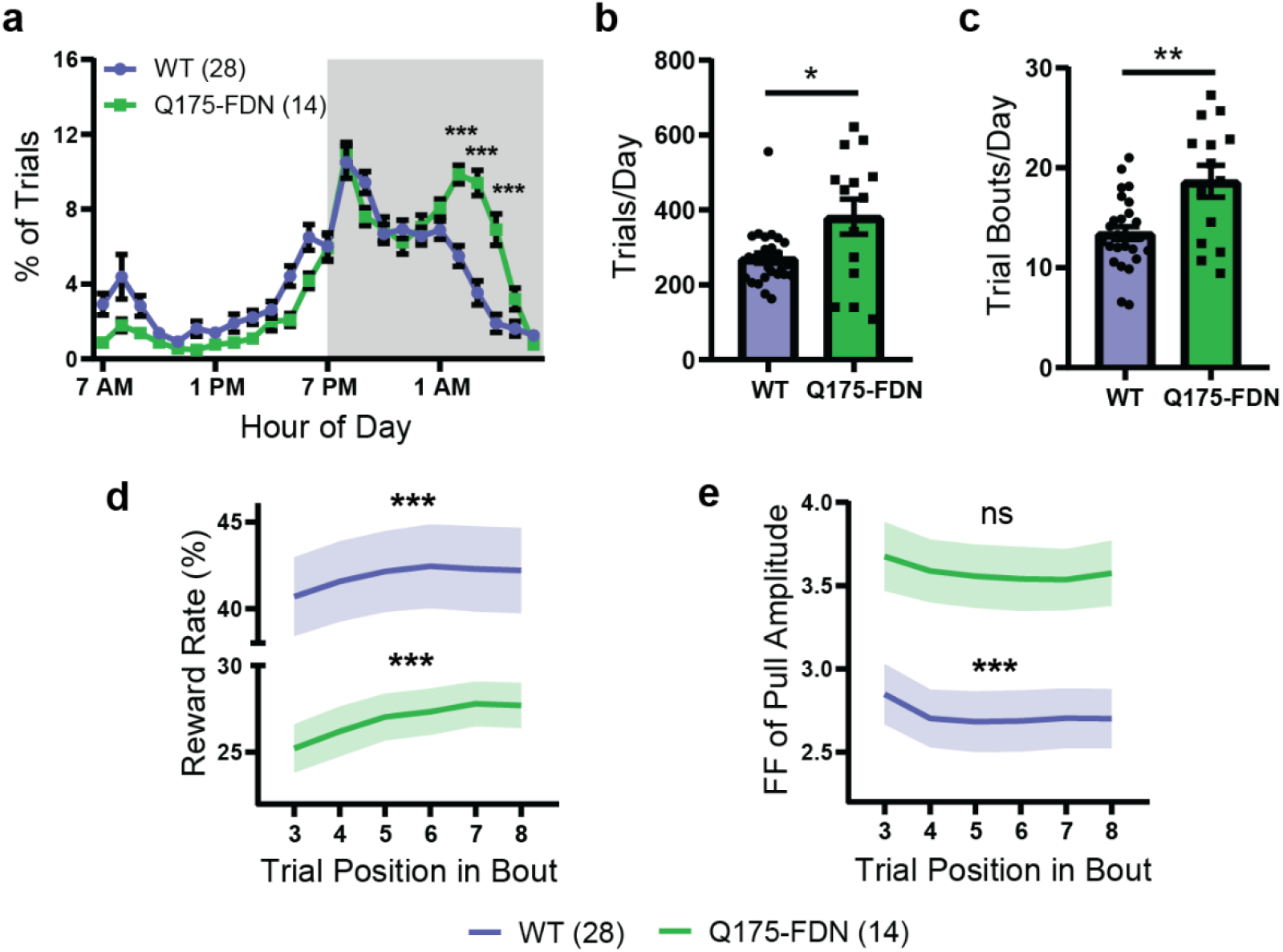
Q175-FDN mice have altered circadian activity levels and impaired short-term motor learning. **(a)** The average circadian distribution of trials (hourly bins) was significantly different in Q175-FDN and WT mice (*p* < 0.0001). Grey-shaded region indicates the dark phase of the light cycle (7 PM – 7 AM). **(b)** Q175-FDN (*n* = 14) mice performed significantly more trials per day as compared to WT mice (*n* = 28). **(c)** Q175-FDN mice (*n* = 14) had a greater number of trial bouts per day as compared to WT mice (*n* = 28). **(d)** Reward rate increased over the course of a bout in both WT (*p* = 0.0004) and Q175-FDN mice (*p* = 0.0006), although it was significantly lower in Q175-FDN overall (*p* = 0.0001). Reward rate was calculated for a 5-trial window centered on the indicated trial position (e.g. trials 1-5 for trial 3). **(e)** The Fano factor of pull amplitude decreased over the course of a bout in WT (*p* = 0.0002), but not Q175-FDN mice (*p* = 0.26) and was significantly higher in Q175-FDN mice overall (*p* = 0.005). Fano factor was calculated for a 5-trial window centered on the indicated trial position. All data are from mice tested on the amplitude task and are presented as mean ± SEM.

To look at within-bout learning, the reward rate was calculated in a 5-trial moving window over each bout with at least 10 trials (as previously). A slight, but significant, increase in average reward rate was seen over the course of each bout in both WT and Q175-FDN mice, although reward rate was overall much lower in the HD group (Trial Position: *F*_5, 200_ = 9.67, *p* < 0.0001; Genotype: *F*_1, 40_ = 18.49, *p* = 0.0001; Interaction: *F*_5, 200_ = 0.63, *p* = 0.68; ANOVA) (Fig. 6d). Conversely, the Fano factor of pull amplitude (5-trial window) was higher in Q175-FDN as compared to WT mice when assessed on the level of individual bouts (Trial Position: *F*_5, 200_ = 5.11, *p* = 0.0002; Genotype: *F*_1, 40_ = 8.93, *p* = 0.0048; Interaction: *F*_5, 200_ = 0.24, *p* = 0.94; ANOVA) (Fig. 6e). Interestingly, although amplitude variability decreased over the course of each bout in WT mice (*F*_5, 135_ = 5.34, *p* = 0.0002, ANOVA), no significant change was seen in the Q175-FDN group (*F*_5, 65_ = 1.34, *p* = 0.26, ANOVA) (Fig. 6e). This suggests that although Q175-FDN mice are able to show some within-session improvements in task performance, these gains are not due to a reduction in movement variability. Overall, these results indicate that on both short and long time-scales, Q175-FDN mice are unable to effectively decrease the variability of their movements in order to increase reward.

### Q175-FDN HD mice have altered learning-associated changes in dorsolateral striatum medium spiny neuronal activity

Motor learning and refinement of movement variability are known to be associated with plasticity of cortico-striatal circuits (Yin et al., 2009; Santos et al., 2015; Kupferschmidt et al., 2017; Giordano et al., 2018). As previous studies have reported altered activity and plasticity of striatal medium spiny neurons (MSNs) in Q175-FDN mice (Southwell et al., 2016; Sepers et al., 2018), one possibility is that the motor learning deficits observed in this model are related to dysfunctional cortico-striatal plasticity. To investigate this, we next used acute slice electrophysiology to measure the spontaneous activity of dorsolateral striatum MSNs in animals who were either task-naïve or had been tested on the PiPaw task. In naïve animals, the frequency of spontaneous excitatory postsynaptic currents (sEPSCs) was markedly higher in WT as compared to Q175-FDN mice (*t*_51_ = 4.93, *p* < 0.0001, t-test; *n* = 22(4) WT, 31(5) Q175-FDN) (Fig. 7a). However, in animals trained on the task, this difference in frequency was no longer significant (*t*_93_ = 1.55, *p* = 0.13, t-test; *n* = 53(8) WT, 42(5) Q175-FDN). In contrast, no genotype differences were observed in sEPSC amplitude in either naïve (*t*_51_ = 1.15, *p* = 0.26, t-test; *n* = 22(4) WT, 31(5) Q175-FDN) or trained animals (*t*_93_ = 1.81, *p* = 0.07, t-test; *n* = 53(8) WT, 42(5) Q175-FDN) (Fig. 7b). The paired-pulse ratio (PPR), a measure of presynaptic probability of release, was found to be significantly higher in Q175-FDN as compared to WT mice (Interval: *F*_4, 256_ = 36.5, *p* < 0.0001; Genotype: *F*_1, 64_ = 0.94, *p* = 0.34; Interaction: *F*_4, 256_ = 4.31, *p* = 0.002; ANOVA; *n* = 42(7) WT, 24(5) Q175-FDN), specifically at the shortest pulse interval of 50ms (*p* = 0.0009; Sidak) (Fig. 7c). This indicates that presynaptic probability of release was lower in Q175-FDN mice as compared to WT, consistent with the decreased frequency of sEPSCs in this group. Again, however, this difference was not seen in trained animals (Interval: *F*_4, 256_ = 26.6, *p* < 0.0001; Genotype: *F*_1, 64_ = 1.63, *p* = 0.21; Interaction: *F*_4, 256_ = 0.58, *p* = 0.68; ANOVA; *n* = 38(6) WT, 28(5) Q175-FDN) (Fig. 7d).

**Figure 7:**
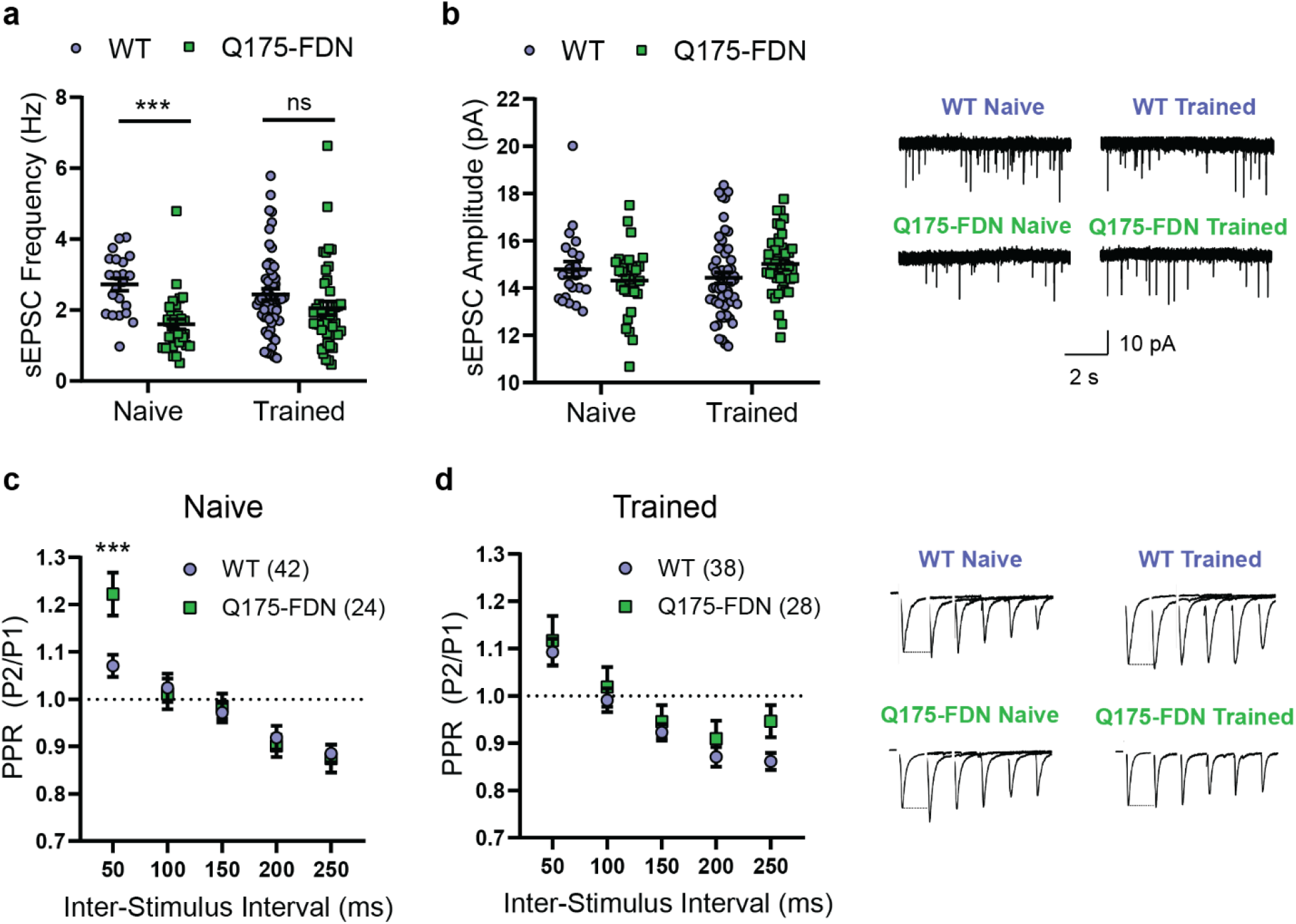
PiPaw training reduces genotype differences in glutamatergic input to dorsolateral striatum medium spiny neurons. **(a)** Spontaneous EPSC (sEPSC) frequency recorded from dorsolateral striatum medium spiny neurons (DLS-MSNs) in 10-month-old WT and Q175-FDN mice who were either trained on the PiPaw task or task-naïve (*n* = 22(4) WT Naive, 31(5) Q175-FDN Naïve, 53(8) WT Trained, 42(5) Q175-FDN Trained). sEPSC frequency is strongly reduced in neurons from naïve Q175-FDN mice, but no genotype differences are seen after training. **(b)** sEPSC amplitude in DLS-MSNs is similar between genotypes and in naïve vs. PiPaw-trained mice (*n* = 22(4) WT Naive, 31(5) Q175-FDN Naïve, 53(8) WT Trained, 42(5) Q175-FDN Trained). Representative traces are shown in right panel. **(c)** Paired-pulse ratio (PPR) recorded from DLS-MSNs in task-naïve WT and Q175-FDN mice (*n* = 42(7) WT, 24(5) Q175-FDN). PPR is significantly higher in Q175-FDN mice at the shortest inter-stimulus interval, indicating a lower presynaptic probability of release. **(d)** PPR is not different in DLS-MSNs from PiPaw-trained WT and Q175-FDN mice (*n* = 38(6) WT, 28(5) Q175-FDN). Representative traces are shown in right panel (stimulation artifact removed for clarity).

Given that training on the PiPaw task is unilateral (i.e. only the right forelimb performs the task), learning-associated changes in the activity of DLS-MSNs are likely to be hemisphere-specific. Thus, we next analyzed patch clamp recordings of MSNs specifically in the left or right DLS to determine if there were hemispheric differences in the spontaneous activity of these neurons. In task-naïve WT and Q175-FDN mice, no hemispheric differences were seen in either sEPSC frequency (Hemisphere: *F*_1, 7_ = 0.44, *p* = 0.53; Genotype: *F*_1, 7_ = 7.78, *p* = 0.03; Interaction: *F*_1, 7_ = 0.01, *p* = 0.91; ANOVA; *n* = 4 WT, 5 Q175-FDN) or amplitude (Hemisphere: *F*_1, 7_ = 0.04, *p* = 0.85; Genotype: *F*_1, 7_ = 0.03, *p* = 0.86; Interaction: *F*_1, 7_ = 2.42, *p* = 0.16; ANOVA; *n* = 4 WT, 5 Q175-FDN) (data not shown). Likewise, no hemispheric differences in sEPSC frequency were seen in animals that had been trained on the task (Hemisphere: *F*_1, 11_ = 0.83, *p* = 0.38; Genotype: *F*_1, 11_ = 1.07, *p* = 0.32; Interaction: *F*_1, 11_ = 0.61, *p* = 0.45; ANOVA; *n* = 8 WT, 5 Q175-FDN) (Fig. 8a). However, a significant effect of hemisphere was observed in the amplitude of sEPSCs in trained mice (Hemisphere: *F*_1, 11_ = 5.5, *p* = 0.039; Genotype: *F*_1, 11_ = 0.86, *p* = 0.37; Interaction: *F*_1, 11_ = 3.26, *p* = 0.1; ANOVA; *n* = 8 WT, 5 Q175-FDN) (Fig. 8b). In WT animals, sEPSC amplitude was consistently lower in the left hemisphere (contralateral to the trained forelimb) as compared to the right hemisphere (*p* = 0.013; Sidak) (Fig. 8b). In contrast, no hemispheric difference was seen in sEPSC amplitude in PiPaw-trained Q175-FDN mice (*p* = 0.93; Sidak) (Fig. 8b).

**Figure 8:**
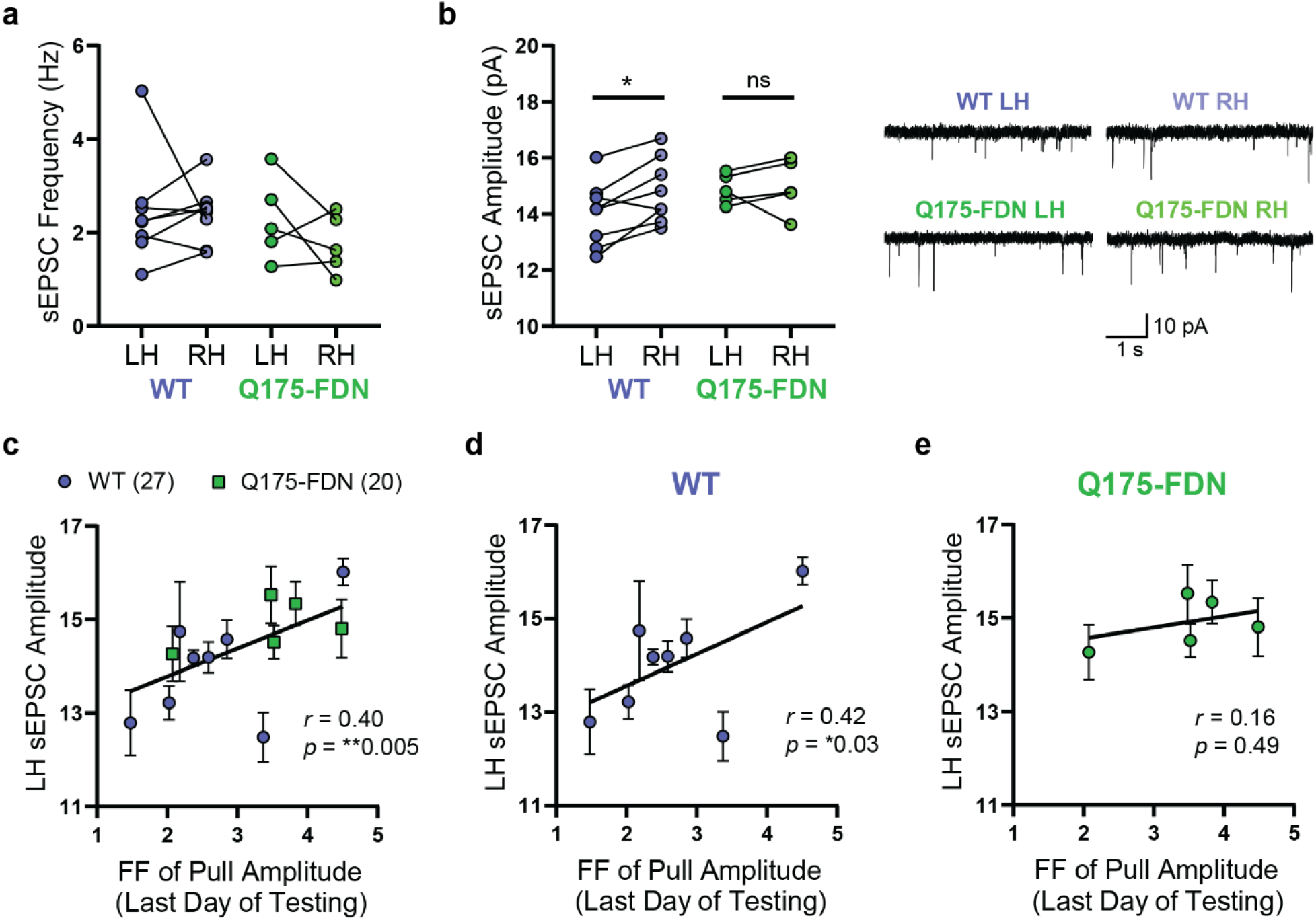
PiPaw training is associated with a decrease in average DLS-MSN sEPSC amplitude in the hemisphere contralateral to the trained forelimb in WT, but not Q175-FDN mice. **(a)** The average frequency of sEPSCs in DLS-MSNs was not significantly different between left and right hemisphere in PiPaw-trained WT and Q175-FDN mice (*n* = 8 WT, 5 Q175-FDN). Lines indicate paired values for each mouse. **(b)** Average sEPSC amplitude in DLS-MSNs was significantly lower in the left hemisphere (contralateral to the trained forelimb) as compared to the right hemisphere in WT animals (*n* = 8), but no hemispheric differences were seen in Q175-FDN mice (*n* = 5). Lines indicate paired values for each mouse. Representative traces are shown in the right panel. **(c)** Average sEPSC amplitude in the left hemisphere (contralateral to the trained forelimb) was significantly correlated with the Fano factor of pull amplitude on the last day of testing across WT and Q175-FDN mice (*n* = 13 mice, 47 cells). **(d)** LH sEPSC amplitude was correlated with the Fano factor of pull amplitude on the last day of testing in WT mice (*n* = 8 mice, 27 cells). **(e)** LH sEPSC amplitude was not correlated with the Fano factor of pull amplitude on the last day of testing in Q175-FDN mice (*n* = 5 mice, 20 cells).

The presence of a consistent hemispheric difference in sEPSC amplitude following PiPaw testing in WT mice suggested that this change in spontaneous neuronal activity was directly related to learning of the task. To further investigate this relationship, we next assessed whether left hemisphere sEPSC amplitude was correlated with any measures of task performance on the last day of testing (immediately prior to electrophysiology experiments). Surprisingly, we found that left hemisphere DLS-MSN sEPSC amplitude was correlated with the Fano factor of pull amplitude on the last day of testing (*r* = 0.4, *p* = 0.005, Pearson; *n* = 47), with those animals showing the least movement variability tending to have the lowest average sEPSC amplitude (Fig. 8c). Separating the two genotypes, we found that this correlation was only significant in WT mice (*r* = 0.42, *p* = 0.03, Pearson; *n* = 27) (Fig. 8d) and was not present in Q175-FDN (*r* = 0.16, *p* = 0.49, Pearson; *n* = 20) (Fig. 8e). These results suggest that in WT animals, task-related refinement of kinematic variability is associated with a decrease in the average amplitude of glutamatergic signals in dorsolateral striatum MSNs. In Q175-FDN mice however, no such relationship is observed, suggesting that learning-associated plasticity of cortico-striatal circuits is impaired.

## Discussion

In both HD patients and pre-symptomatic carriers of the HTT mutation, various deficits have been found in the ability to learn and control voluntary movements. For example, HD patients have impaired learning of motor sequence tasks (Willingham and Koroshetz, 1993; Feigin et al., 2006), and reaching movements display jerkiness, impaired error correction and abnormal temporal sequencing of movement steps (Bonfiglioli et al., 1998; Smith et al., 2000; Klein et al., 2011; Shabbott et al., 2013). Although tests of skilled forelimb use in rodents have good face and construct validity for the functional assessment of striatal pathology (Karl and Whishaw, 2011; Klein et al., 2012), they have seen limited use in mouse models of HD. Here, we found that Q175-FDN HD mice were significantly impaired at performing the PiPaw task, an operant forelimb motor learning assessment that is integrated into the mouse home-cage. Despite having similar initial performance to wildtype animals, Q175-FDN mice were unable to effectively decrease the variability of their movements in order to increase their reward rate on either short (minutes) or long (days) timescales.

In wildtype mice, improved performance of the task was primarily driven by a decrease in the inter-trial variability of pull kinematics across days of testing. Variability during the early stages of reinforcement learning is advantageous, as exploration is necessary in order to determine execution parameters that will lead to reward (Dhawale et al., 2017; Van Mastrigt et al., 2020). However, as information about the outcomes of different movements becomes available, variability is reduced in order to improve performance (Pekny et al., 2015; Dhawale et al., 2019). As movement execution is inherently noisy and it may not be possible to decrease variability universally, an optimal solution is to reduce variability preferentially in parameters that are relevant to task success (Todorov and Jordan, 2002; Santos et al., 2015). In this case, the inter-trial variability of pull amplitude, the parameter most relevant to reward, decreased by an average of 42% over one week of testing. In addition, amplitude variability was highly inversely correlated with daily reward rate across animals, suggesting that mice bidirectionally modulate the variability of this parameter in response to success or failure. In a separate paradigm in which reward delivery was instead contingent on pull velocity, the inter-trial variability of peak velocity decreased with increasing reward rate, while the variability of pull amplitude increased. Together, these results indicate that mice tested on this task were sensitive to differing reward contingencies and independently regulated the trial-to-trial variability of specific kinematic parameters in order to increase reward. Interestingly, the motor learning deficit in Q175-FDN animals was not due to a failure to reduce the variability of their movements *per se*, as the Fano factor of pull amplitude was found to decrease over time on a group level. However, this decrease was not correlated with improved performance of the task as it was in WT mice. This suggests that the decrease in variability was non-specific, rather than a response to an increase or decrease in reward rate.

One of the most compelling benefits of studying behaviour in a home-cage system is the opportunity to examine the structure of self-paced learning and task performance. In this regard, we found that mice consistently clustered their trials into short bouts of high task engagement, followed by longer breaks where no trials were performed. The large majority of trials occurred in such bouts, with less than 10% of trials being classified as non-bout trials. On one hand, this structure is likely related to the natural circadian pattern of eating and drinking displayed by mice (Ho and Chin, 1988; Godynyuk et al., 2019) and the volume of water required to sate the animals thirst. However, there also appeared to be a more functional purpose of clustering trials into short bouts. Animals displayed significant within-bout learning, with trials occurring later in a bout having a higher reward rate and lower amplitude variability than those early in the bout. This short-term learning is often seen in motor tasks and tends to be most prominent in early stages of learning (Buitrago et al., 2004a, 2004b). Although Q175-FDN mice also showed within-bout improvements in reward rate, this was not due to changes in inter-trial variability of pull amplitude. Instead, short-term improvements in performance may have been driven by changes in the average amplitude of pulls (e.g. towards the center of the rewarded range), or by a decrease in the number of pulls held for longer than the trial time limit. This pattern of slight improvements in task performance that are unrelated to decreased movement variability recapitulates the general trend observed in Q175-FDN mice and suggests that these animals employed alternative strategies to improve their reward rate.

Surprisingly, YAC128 HD mice at a timepoint when they exhibit movement abnormalities showed intact motor learning and refinement of kinematic variability on the PiPaw task. The divergent phenotype between these two mouse models of HD was surprising given that YAC128 mice are reported to manifest motor abnormalities several months earlier than Q175-FDN mice (Slow et al., 2003; Southwell et al., 2016). However, previous studies of YAC128 mice have generally relied on tests of full body motor coordination such as the rotarod for defining the onset of motor symptoms. Increased bodyweight, as is seen in the YAC128 model, is known to impair performance on the rotarod test (McFadyen et al., 2003), and restoring normal body weight with dietary restriction was found to rescue rotarod deficits in YAC128 mice (Moreno et al., 2016). Given the low baseline levels of locomotor activity in these mice, fatigue may also play a role in poor rotarod performance, as rotarod testing protocols can involve up to five minutes of continuous locomotion. It is important to note, however, that subtle motor impairments have previously been reported on reaching and lever manipulation tasks in YAC128 mice, specifically when the demands of the task change or when there is a break in testing (Woodard et al., 2017; Glangetas et al., 2020). Still, our results suggest that motor learning deficits in YAC128 mice may not be as severe as previously thought and encourage further elucidation of the motor phenotype in these mice.

A number of studies have found that motor learning is associated with changes in the activity of cortical inputs to the DLS (Kupferschmidt et al., 2017) and DLS-MSNs themselves (Yin et al., 2009; Santos et al., 2015; Shan et al., 2015; Giordano et al., 2018). In wildtype mice, we found that training on the PiPaw task was associated with a decrease in the amplitude of spontaneous excitatory events specifically in DLS-MSNs contralateral to the trained forelimb. Furthermore, sEPSC amplitude was positively correlated with the inter-trial variability of pull amplitude in each animal, suggesting that the observed postsynaptic depression was related to increased movement stereotypy. In contrast, no electrophysiological hemispheric differences were observed in Q175-FDN mice following PiPaw-training, and DLS-MSN sEPSC amplitude was not correlated with pull amplitude variability in this group. The absence of any change in the strength of glutamatergic synapses onto striatal MSNs in Q175-FDN mice in response to training on this task is consistent with previous findings of impaired long-term potentiation (LTP) and long-term depression (LTD) in this mouse model (Sepers et al., 2018; Quirion and Parsons, 2019). Indeed, given that mouse models of HD have comparatively minimal neuronal degeneration overall, behavioural deficits in these mice may be primarily due to synaptic and circuit level dysfunction (Raymond et al., 2011; Plotkin and Surmeier, 2015). Thus, the presence of a behavioural phenotype that directly correlates with a neurophysiological measure is promising and encourages further use of this system for the assessment of HD mice.

Home-cage testing systems offer several important advantages over more traditional behavioural paradigms. In addition to decreasing the exposure of animals to handling and other potential stressors, automated home-cage systems greatly increase the throughput of behavioural experiments. Over the course of testing (1 to 4 weeks), several thousand trials were collected for each mouse with minimal experimenter intervention aside from standard animal husbandry. Overall, the PiPaw system exhibited a high success rate in training both young and aged mice, and animals generally responded well to the shaping paradigm and changes in task reward requirements. We were able to measure motor kinematics with high temporal (~5 ms) and position (~50 μm) resolution, allowing us to assess motor learning in HD mice with an unprecedented level of detail. In addition, the group housing accommodated by this system allowed for heterozygous transgenic or knock-in mice to be tested alongside their WT littermates, providing within-cage control of environmental factors. These results further validate the use of this system (and home-cage tools more generally) for behavioural assessment in genetic models of Huntington disease and other neurological disorders.

## Acknowledgments

This study was supported by the Canadian Institutes of Health Research (CIHR) Grant FDN-143210 to LAR. CLW holds a CIHR Canada Graduate Scholarship-Doctoral. The authors are grateful to Dr. Michael Hayden, University of British Columbia, for providing the Q175-FDN mouse line used in these experiments. We are also grateful to Luis Bolaños for his assistance with figures and to Lily Zhang for animal care support and genotyping.

## Notes

Conflict of Interest: The authors declare no competing financial interests.

### Competing Interest Statement

The authors have declared no competing interest.

## References

Abada Y-SK, Schreiber R, Ellenbroek B (2013) Motor, emotional and cognitive deficits in adult BACHD mice: a model for Huntington’s disease. Behav Brain Res 238:243–251.

Balcombe JP, Barnard ND, Sandusky C (2004) Laboratory routines cause animal stress. J Am Assoc Lab Anim Sci 43:42–51.

Bates GP, Dorsey R, Gusella JF, Hayden MR, Kay C, Leavitt BR, Nance M, Ross CA, Scahill RI, Wetzel R, Wild EJ, Tabrizi SJ (2015) Huntington disease. Nat Rev Dis Prim 1:1–21.

Bollu T, Whitehead SC, Prasad N, Walker J, Shyamkumar N, Subramaniam R, Kardon B, Cohen I, Goldberg JH (2019) Automated home cage training of mice in a hold-still center-out reach task. J Neurophysiol 121:500–512.

Bonfiglioli C, Berti G De, Nichelli P, Nicoletti R, Castiello U (1998) Kinematic analysis of the reach to grasp movement in Parkinson’s and Huntington’s disease subjects. Neuropsychologia 36:1203–1208.

Buitrago MM, Ringer T, Schulz JB, Dichgans J, Luft AR (2004a) Characterization of motor skill and instrumental learning time scales in a skilled reaching task in rat. Behav Brain Res 155:249–256.

Buitrago MM, Schulz JB, Dichgans J, Luft AR (2004b) Short and long-term motor skill learning in an accelerated rotarod training paradigm. Neurobiol Learn Mem 81:211–216.

Dhawale AK, Miyamoto YR, Smith MA, Ölveczky BP (2019) Adaptive regulation of motor variability. Curr Biol 29:3551–3562.

Dhawale AK, Smith MA, Ölveczky BP (2017) The role of variability in motor learning. Annu Rev Neurosci 40:479–498.

Feigin A, Ghilardi MF, Huang C, Ma Y, Carbon M, Guttman M, Paulsen JS, Ghez CP, Eidelberg D (2006) Preclinical Huntington’s disease: compensatory brain responses during learning. Ann Neurol 59:53–59.

Giordano N, Iemolo A, Mancini M, Cacace F, De Risi M, Latagliata EC, Ghiglieri V, Bellenchi GC, Puglisi-Allegra S, Calabresi P, Picconi B, De Leonibus E (2018) Motor learning and metaplasticity in striatal neurons: relevance for Parkinson’s disease. Brain 141:505–520.

Glangetas C, Espinosa P, Bellone C (2020) Deficit in Motor Skill Consolidation-Dependent Synaptic Plasticity at Motor Cortex to Dorsolateral Striatum Synapses in a Mouse Model of Huntington’s Disease. eNeuro 7.

Godynyuk E, Bluitt MN, Tooley JR, Kravitz A V, Creed MC (2019) An open-source, automated home-cage sipper device for monitoring liquid ingestive behavior in rodents. eNeuro 6.

Graybiel AM, Grafton ST (2015) The striatum: where skills and habits meet. Cold Spring Harb Perspect Biol 7.

Ho A, Chin A (1988) Circadian feeding and drinking patterns of genetically obese mice fed solid chow diet. Physiol Behav 43:651–656.

Karl JM, Whishaw IQ (2011) Rodent skilled reaching for modeling pathological conditions of the human motor system. In: Neuromethods (Lane EL, Dunnett SB, eds), pp 87–107. Humana Press.

Klaus A, Alves Da Silva J, Costa RM (2019) What, if, and when to move: basal ganglia circuits and self-paced action initiation. Annu Rev Neurosci 42:459–483.

Klein A, Sacrey LAR, Dunnett SB, Whishaw IQ, Nikkhah G (2011) Proximal movements compensate for distal forelimb movement impairments in a reach-to-eat task in Huntington’s disease: new insights into motor impairments in a real-world skill. Neurobiol Dis 41:560–569.

Klein A, Sacrey LR, Whishaw IQ, Dunnett SB (2012) The use of rodent skilled reaching as a translational model for investigating brain damage and disease. Neurosci Biobehav Rev 36:1030–1042.

Kudo T, Schroeder A, Loh DH, Kuljis D, Jordan MC, Roos KP, Colwell CS (2011) Dysfunctions in circadian behavior and physiology in mouse models of Huntington’s disease. Exp Neurol 228:80–90.

Kupferschmidt DA, Juczewski K, Cui G, Johnson KA, Lovinger DM (2017) Parallel, but dissociable, processing in discrete corticostriatal inputs encodes skill learning. Neuron 96:476–489.

Loh DH, Kudo T, Truong D, Wu Y, Colwell CS (2013) The Q175 mouse model of Huntington’s disease shows gene dosage- and age-related decline in circadian rhythms of activity and sleep. PLoS One 8.

McColgan P, Tabrizi SJ (2018) Huntington’s disease: a clinical review. Eur J Neurol 25:24–34.

McFadyen MP, Kusek G, Bolivar VJ, Flaherty L (2003) Differences among eight inbred strains of mice in motor ability and motor learning on a rotorod. Genes, Brain Behav 2:214–219.

Moreno CL, Ehrlich ME, Mobbs C V. (2016) Protection by dietary restriction in the YAC128 mouse model of Huntington’s disease: relation to genes regulating histone acetylation and HTT. Neurobiol Dis 85:25–34.

Morton AJ, Wood NI, Hastings MH, Hurelbrink C, Barker RA, Maywood ES (2005) Disintegration of the sleep-wake cycle and circadian timing in Huntington’s disease. J Neurosci 25:157–163.

Murphy TH, Boyd JD, Bolaños F, Vanni MP, Silasi G, Haupt D, Ledue JM (2016) High-throughput automated home-cage mesoscopic functional imaging of mouse cortex. Nat Commun 7.

Pekny SE, Izawa J, Shadmehr R (2015) Reward-dependent modulation of movement variability. J Neurosci 35:4015–4024.

Plotkin JL, Surmeier DJ (2015) Corticostriatal synaptic adaptations in Huntington’s disease. Curr Opin Neurobiol 33:53–62.

Poddar R, Kawai R, Ölveczky BP (2013) A fully automated high-throughput training system for rodents. PLoS One 8.

Pouladi MA, Morton AJ, Hayden MR (2013) Choosing an animal model for the study of Huntington’s disease. Nat Rev Neurosci 14:708–721.

Pouladi MA, Xie Y, Skotte NH, Ehrnhoefer DE, Graham RK, Kim JE, Bissada N, Yang XW, Paganetti P, Friedlander RM, Leavitt BR, Hayden MR (2010) Full-length huntingtin levels modulate body weight by influencing insulin-like growth factor 1 expression. Hum Mol Genet 19:1528–1538.

Quirion JG, Parsons MP (2019) The onset and progression of hippocampal synaptic plasticity deficits in the Q175FDN mouse model of Huntington disease. Front Cell Neurosci 13.

Raymond LA, André VM, Cepeda C, Gladding CM, Milnerwood AJ, Levine MS (2011) Pathophysiology of Huntington’s disease: time-dependent alterations in synaptic and receptor function. Neuroscience 198:252–273.

Salameh G, Jeffers MS, Wu J, Pitney J, Silasi G (2020) The home-cage automated skilled reaching apparatus (HASRA): Individualized training of group-housed mice in a single pellet reaching task. eNeuro 7.

Santos FJ, Oliveira RF, Jin X, Costa RM (2015) Corticostriatal dynamics encode the refinement of specific behavioral variability during skill learning. Elife 4.

Sepers MD, Smith-Dijak AI, Ledue J, Kolodziejczyk K, Mackie K, Raymond LA (2018) Endocannabinoid-specific impairment in synaptic plasticity in striatum of Huntington’s disease mouse model. J Neurosci 38:544–554.

Shabbott B, Ravindran R, Schumacher JW, Wasserman PB, Marder KS, Mazzoni P (2013) Learning fast accurate movements requires intact frontostriatal circuits. Front Hum Neurosci 7.

Shan Q, Christie MJ, Balleine BW (2015) Plasticity in striatopallidal projection neurons mediates the acquisition of habitual actions. Eur J Neurosci 42:2097–2104.

Silasi G, Boyd JD, Bolanos F, LeDue JM, Scott SH, Murphy TH (2018) Individualized tracking of self-directed motor learning in group-housed mice performing a skilled lever positioning task in the home cage. J Neurophysiol 119:337–346.

Slow EJ, van Raamsdonk J, Rogers D, Coleman SH, Graham RK, Deng Y, Oh R, Bissada N, Hossain SM, Yang YZ, Li XJ, Simpson EM, Gutekunst CA, Leavitt BR, Hayden MR (2003) Selective striatal neuronal loss in a YAC128 mouse model of Huntington disease. Hum Mol Genet 12:1555–1567.

Smith MA, Brandt J, Shadmehr R (2000) Motor disorder in Huntington’s disease begins as a dysfunction in error feedback control. Nature 403:544–549.

Sorge RE et al. (2014) Olfactory exposure to males, including men, causes stress and related analgesia in rodents. Nat Methods 11:629–632.

Southwell AL, Smith-Dijak AI, Kay C, Sepers MD, Villanueva EB, Parsons MP, Xie Y, Anderson L, Felczak B, Waltl S, Ko S, Cheung D, Cengio LD, Slama R, Petoukhov E, Raymond LA, Hayden MR (2016) An enhanced Q175 knock-in mouse model of Huntington disease with higher mutant huntingtin levels and accelerated disease phenotypes. Hum Mol Genet 25:3654–3675.

Todorov E, Jordan MI (2002) Optimal feedback control as a theory of motor coordination. Nat Neurosci 5:1226–1235.

Van Mastrigt NM, Smeets JBJ, Van Der Kooij K (2020) Quantifying exploration in reward-based motor learning. PLoS One 15.

Willingham DB, Koroshetz WJ (1993) Evidence for dissociable motor skills in Huntington’s disease patients. Psychobiology 21:173–182.

Woodard CL, Bolaños F, Boyd JD, Silasi G, Murphy TH, Raymond LA (2017) An automated home-cage system to assess learning and performance of a skilled motor task in a mouse model of Huntington’s disease. eNeuro 4.

Woodard CL, Nasrallah WB, Samiei B V., Murphy TH, Raymond LA (2020) PiDose: an open-source system for accurate and automated oral drug administration to group-housed mice. Sci Rep 10.

Yin HH, Mulcare SP, Hilário MRF, Clouse E, Holloway T, Davis MI, Hansson AC, Lovinger DM, Costa RM (2009) Dynamic reorganization of striatal circuits during the acquisition and consolidation of a skill. Nat Neurosci 12:333–341.

